# MAGIC: A label-free fluorescence method for 3D high-resolution reconstruction of myelinated fibers in large volumes

**DOI:** 10.1101/2020.07.28.225011

**Authors:** Irene Costantini, Enrico Baria, Michele Sorelli, Felix Matuschke, Francesco Giardini, Miriam Menzel, Giacomo Mazzamuto, Ludovico Silvestri, Riccardo Cicchi, Katrin Amunts, Markus Axer, Francesco Saverio Pavone

## Abstract

Analyzing the structure of neuronal fibers with single axon resolution, in large volumes, remains an unresolved challenge in connectomics. Here, we propose MAGIC (Myelin Autofluorescence imaging by Glycerol Induced Contrast enhancement), a simple tissue preparation method to perform label-free fluorescence imaging of myelinated fibers. We demonstrate its broad applicability by performing mesoscopic reconstruction at sub-micron resolution of mouse, rat, monkey, and human brain samples and by quantifying the different fiber organization in Control and Reeler mouse’s hippocampal sections.

## Main

The brain is a complex organ constituted by highly interconnected units, the neurons, capable of storing and processing information from a myriad of different inputs regulating most human activities. The aim of connectomics consists in reconstructing the intricate organization of the connections between brain regions at macro- to mesoscales, but also between individual neurons at the microscale. Most long-range projecting axons are wrapped by a myelin sheath to permit a reliable and efficient signal transmission^1^. Several methods have been developed to map the network of interneuronal connections; still, they are limited by the volume that they can analyze (Electron Microscopy)^2^, by the spatial resolution achievable (MRI)^3^ or by the sophisticated equipment needed for the measurement (CARS, THG microscopy)^4^. Optical methods have the potential for scalable large-area high-resolution mapping. However, they need a source of contrast to detect the structure of interest. In this respect, myelin staining is an unmet technical challenge. Exogenous dyes^5^ are used to stain fibers composing the white matter, but nonspecific binding and inefficient diffusion of dyes hinder single fiber imaging in large volumes.

To meet this need, we develop MAGIC (Myelin Autofluorescence imaging by Glycerol Induced Contrast enhancement), a simple label-free method that opens the possibility of performing sub-micron resolution fluorescence imaging of myelinated fibers in 3D at the mesoscale level. MAGIC is a methodology that enables, with a glycerol-based procedure, to enhance myelin’s autofluorescence, allowing the use of conventional fluorescence microscopy techniques to investigate neuronal filament organization. Glycerol has been widely used as a mounting and refractive index matching medium because of its biocompatibility^6^. We implemented the MAGIC protocol from the observation that the removal of glycerol from previously fixed and embedded tissue allows for the specific enhancement of myelin autofluorescence. MAGIC includes three steps (Figure 1a): fixation with paraformaldehyde (PFA), embedding in glycerol (Gly), and removal of glycerol by washing in saline solution (MAGIC). During the procedure, myelinated fibers undergo a specific increase of fluorescence efficiency, allowing for high-resolution 3D reconstruction of the axons (Figure 1b, Supplementary Video 1). The number of emitted photons rises significantly during the different steps of the protocol (Figure 1c). In PFA, we observed a negative contrast between fibers and surrounding tissue fluorescence; instead, after the MAGIC protocol, the contrast becomes positive and is significantly increased due to the raising of myelinated fiber autofluorescence (Figure 1d). The protocol does not introduce any exogenous fluorophores and is based on the local enhancement of autofluorescence from endogenous molecules. The fluorescence emitted from the myelinated fibers can be detected not only with two-photon excitation but also with conventional one-photon microscopy. Supplementary figures display images acquired with a commercial confocal microscope at various wavelengths (Supplementary Figure 1) and from various species (Supplementary Figure 2). We demonstrate that the MAGIC protocol can provide details of myelin substructures (Supplementary Figure 3) using a high-magnification objective and that it is also compatible with conventional immunofluorescence (Supplementary Figure 4, Supplementary Video 2 and 3).

**Figure 1.**
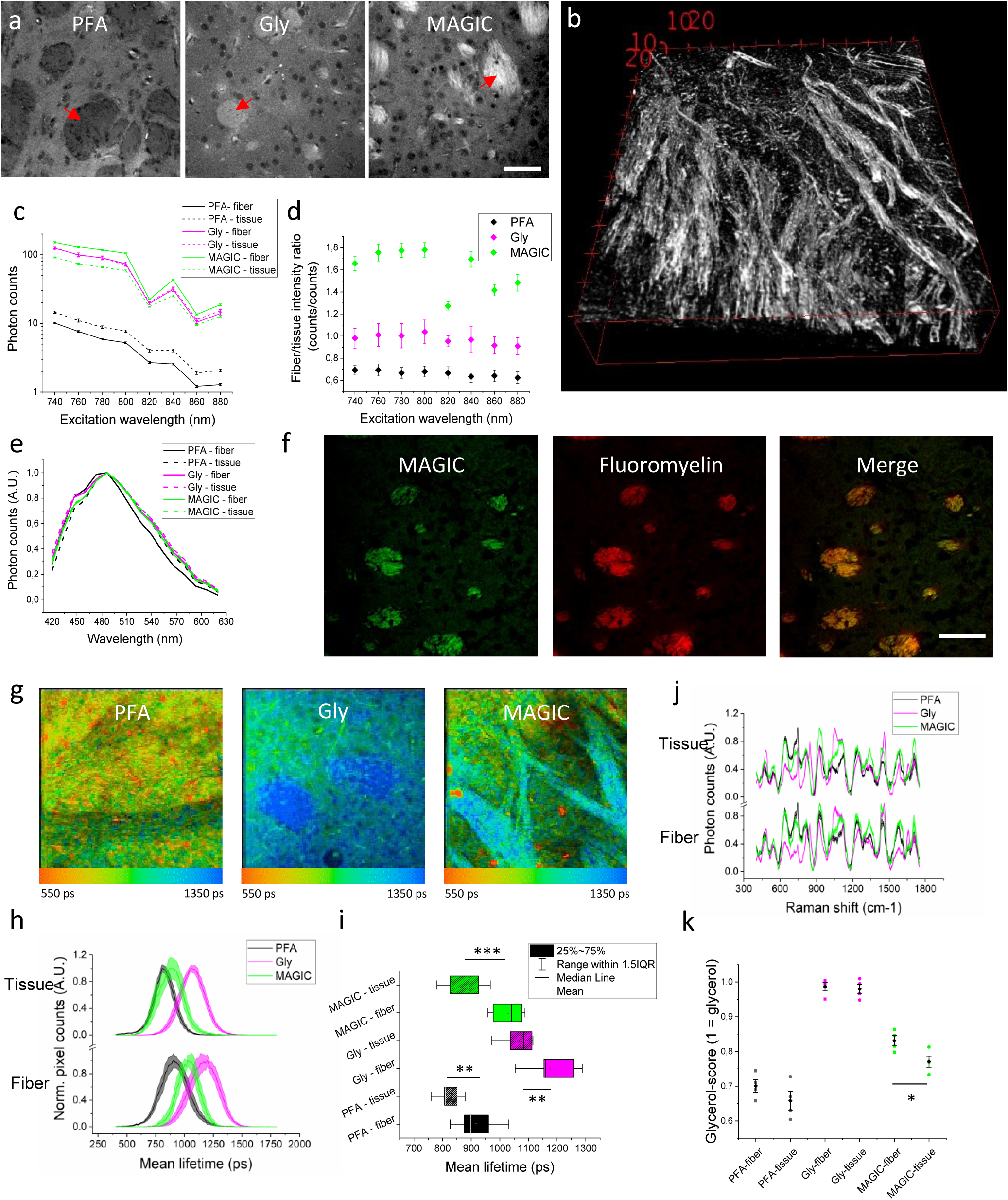
MAGIC: a method to perform label-free fluorescence imaging of myelinated fibers. (a) Representative TPFM images of the caudate putamen of a mouse brain section during the three subsequent steps of the MAGIC protocol: fixation (PFA), glycerolization (Gly), and washing (MAGIC). Red arrows indicate fiber bundles. Scale bar = 50 μm. (b) 3D reconstruction of myelinated fibers, imaging performed with TPFM at the resolution of (0.44 × 0.44 × 1) μm^3^. Box scale = 10 μm. (c) Measurement (mean ± std.err) of photons emitted by the myelinated fibers (fiber) and the surrounding tissue (tissue) during the different steps of the protocol (PFA, Gly, MAGIC) detected at different excitation wavelengths. (d) Intensity contrast (mean ± std.err) observed at different excitation wavelengths during the three steps of the MAGIC protocol. (e) Fluorescence emission spectra of the different samples excited with TPFM at 800 nm. (f) Images of a mouse brain section treated with MAGIC and labeled with FluoroMyelin™ red. In green and red respectively, the autofluorescence and the exogenous signals are shown. Images were obtained with TPFM, scale bar = 50 μm. (g) FLIM representative images during the three steps of the MAGIC protocol. Lifetime color scale is set from 550 to 1350 ps. (h) Fluorescence lifetime distributions (mean ± std.err) of fiber and tissue during the MAGIC steps. (i) Box chart plot of fluorescence lifetime values. (j) Raman spectra (mean ± std.err) of fiber and tissue during the MAGIC steps. (k) Glycerol scores (mean ± std.err) calculated along with the three major bands (550, 850, and 1465 cm^-1^) of the glycerol Raman spectrum. Statistical significance (t-test): *p < 0.05, ** p < 0.001, *** p < 0.0001

**Figure 2.**
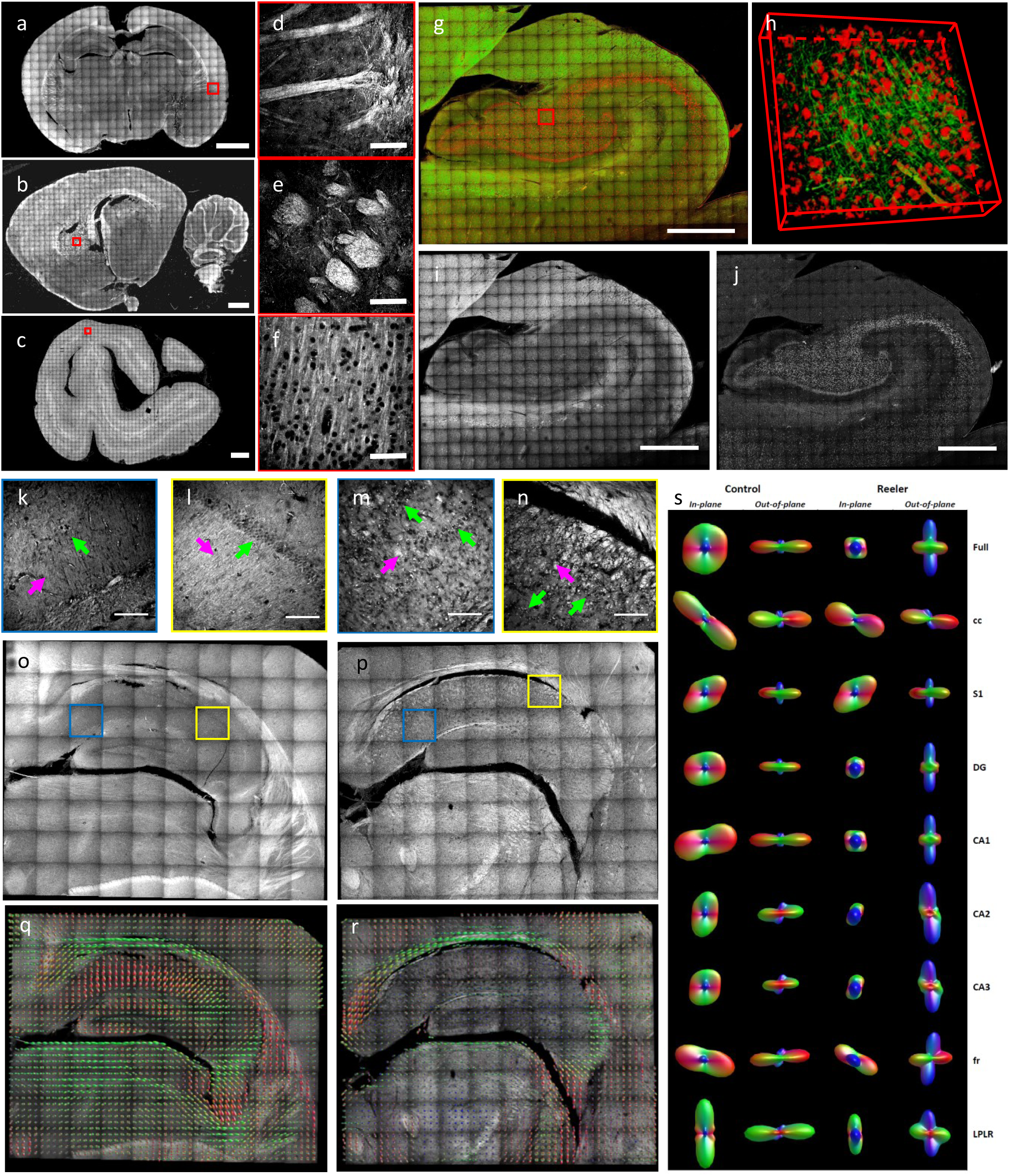
Mesoscopic 3D reconstruction and quantification with MAGIC. Maximum intensity projection (MIP) of the mesoscale reconstruction of 60-µm-thick brain sections treated with MAGIC: mouse (a), rat (b), and vervet monkey (c), respectively. Scale bar = 1 mm. (d, e, f) Magnified inset corresponding respectively to the red boxes in a, b, c. Scale bar = 50 μm. (g) MIP of the mesoscale reconstruction of a human hippocampus 60-µm-thick coronal section. Scale bar = 1 mm. (h) 3D rendering (450 × 450 x 60 µm^3^) of the stack indicated by the red box in g. (i) Green channel showing the myelinated fibers enhanced by MAGIC. (j) The red channel of the MIP in g showing the cells’ bodies autofluorescence produced by lipofuscin pigments. (k, l, m, n) Magnified inset of blue and yellow boxes in q and r, respectively. Green arrows point to neuronal cell bodies, magenta arrows to myelinated fibers. Scale bar = 50 μm. (o, p) MIP of the mesoscale reconstructions of a control and Reeler mouse hippocampus 60-µm-thick coronal section. Scale bar = 1 mm. (q, r) Images show the ODF maps obtained from analyzing the full 3D hippocampus reconstruction of the Control and the Reeler mouse, sampling 16 vectors for each ODF. (s) In-plane and out-of-plane orientation of the single ODF obtained by analyzing all the vectors of the full mosaic reconstructions and each of its ROIs. Acronyms list = Full: full field of view; cc: Corpus Callosum; S1: Primary Somatosensory Cortex; DG: Dentate Gyrus; CA1, CA2, CA3: field CA1, CA2, CA3 of hippocampus; fr: Fasciculus Retroflexus; LPLR: Lateral Posterior Thalamic Nucleus.

The spectral analysis of fluorescence signals reveals that the emission spectrum is not altered throughout the steps of the protocol (Figure 1e). Nevertheless, the fluorescence signal from myelinated fibers is specifically enhanced by MAGIC, as proved by the correlation with a mouse brain section labeled with an exogenous dye specific for myelin staining: FluoroMyelin™ red^7^. The signal emitted by the dye perfectly overlaps the fibers’ autofluorescence signal, indicating that the fluorescence is indeed coming from myelin (Figure 1f).

To investigate the origin of the phenomenon, we analyzed the optical properties of mouse brain sections during the three steps of the protocol. Time-resolved analysis of the emitted photons, obtained with FLIM (Fluorescence-lifetime imaging microscopy)^8^, highlights a fluorescence lifetime increase after glycerolization. MAGIC modifies the dynamics of the fluorescence decay originating from fibers and tissue. The fluorescence lifetime of the tissue decreases at values comparable to those ones obtained from PFA samples (with only 60±30 ps average difference), while that of the fibers remains at significantly higher values, resulting in a 110±40 ps difference from the corresponding PFA lifetime distribution (Figure 1g, 1h, and 1i). These findings suggest that the change of the molecular environment surrounding the fluorescent molecules occurs differently inside and outside the fibers. To further characterize this phenomenon, we used Raman spectroscopy^9^ as a tool to probe molecular content and to prove the involvement of glycerol in the fluorescence emission enhancement of myelinated fibers (Figure 1j). Glycerolized tissue spectra are characterized by the addition of glycerol Raman peaks around 485, 550, 850, 925, 1060, and 1465 cm^-1^ (Supplementary Figure 5a), as compared to PFA samples; such spectral signatures typical of glycerol disappear in the surrounding tissue after MAGIC, whereas they are preserved within myelinated fibers. Conversely, no DMSO contributions (present in the first step of the protocol to enhance the penetration of glycerol) were detected from the Raman spectra (Supplementary Figure 5b). A more detailed analysis based on the main Raman bands of glycerol was performed to quantitatively evaluate the involvement of glycerol in the process (Figure 1k). After MAGIC, myelinated fibers show, on average, a significantly smaller decrease (∼16%) in the intensity of glycerol-related Raman peaks with respect to the surrounding tissue (∼21%). All these findings indicate that the glycerol plays a central role in the MAGIC protocol. The higher fluorescence lifetime measured in the fibers suggests an anti-quenching effect. Raman spectroscopy proves the involvement of glycerol in the fluorescence emission of myelinated fibers. In conclusion, the fluorescence enhancement could be due to the glycerol that remains confined inside the myelinated fibers after MAGIC due to its higher affinity^10^ to this structure with respect to the surrounding tissue.

A significant advantage of the MAGIC protocol is its widespread applicability due to the fact that glycerol is a very common biocompatible mounting medium for tissue samples. MAGIC is highly versatile and can be successfully applied to a wide variety of samples. Brain sections from different mammal species: mouse (Figure 2a, 2d), rat (Figure 2b, 2e), vervet monkey (Figure 2c, 2f), and, more importantly, humans (Figure 2g, 2h, 2i, 2j) can be imaged after being treated with the protocol. Interestingly, the presence of lipofuscin pigments^11^ in the human brain sample allowed, in combination with MAGIC, the label-free detection and the 3D reconstruction of neuronal fibers and cell bodies (Figure 2i, 2j, and Supplementary Video 4), obtaining a more comprehensive anatomical organization of the human hippocampus.

Finally, in order to demonstrate that the MAGIC protocol allows us to characterize the 3D tissue anatomy, we compared the structural organization of different regions of the hippocampus from a Control and a Reeler mouse. Reeler mouse (Reelin deficient - RELN-/- Reeler) is a well-known animal model for several neurological and neurodegenerative disorders^12,13^. In the Reeler sample, the organization of neuronal cell bodies and fibers is different compared to that of the Control mouse (Figure 2k, 2l, 2m, 2n). To quantify the observed alteration on the mesoscale reconstruction (Figure 2o, 2p), different regions of the mosaic were selected and manually segmented according to their anatomical classification (Supplementary Figure 6). A custom-made automatic Structure Tensor Analysis^14^ tool followed by an Orientation Distribution Function (ODF)^15^ evaluation of the derived vectors were applied to the full reconstruction and on each ROI (Figure 2q, 2r, 2s, Supplementary Figure 7). We found that the primary peaks orientation of the Control mouse is mostly constituted by in-plane contributions in all the selected ROIs, while six out of eight areas of the Reeler sample are distributed in the out-of-plane direction (Figure 2s and Table 1). These findings demonstrate that the Reeler mouse’s inner connectivity of the hippocampus differs significantly from the fiber organization of a normal mouse.

**Table 1.**
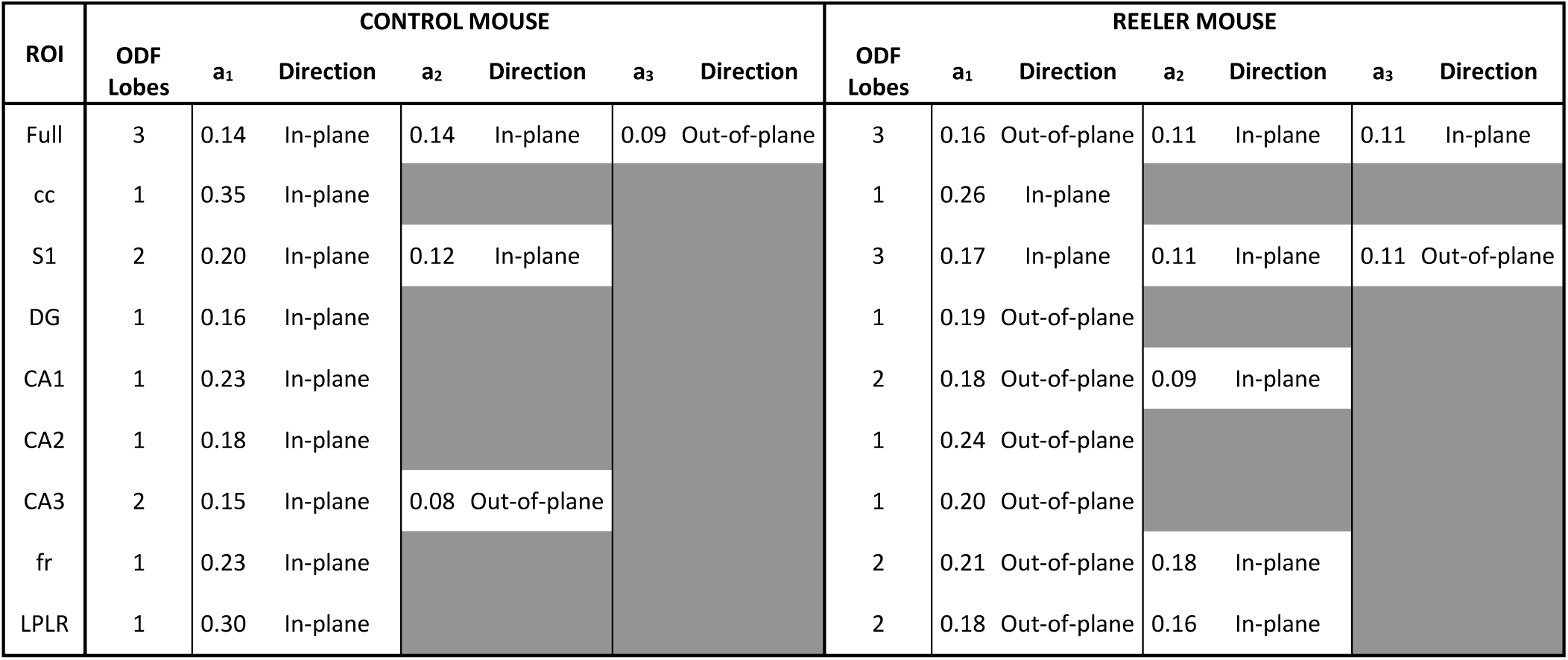
ODF components analysis. Amplitude (a1, a2, a3) and main orientation (direction) of the significant ODF components of both Control and Reeler mouse hippocampi are shown. Table cell is gray if the peak amplitude is <50% of the primary peak. Acronyms list = Full: full field of view; S1: Primary Somatosensory Cortex; cc: Corpus Callosum; CA1, CA2, CA3: field CA1, CA2, CA3 of hippocampus; DG: Dentate Gyrus; fr: Fasciculus Retroflexus; LPLR: Lateral Posterior Thalamic Nucleus.

To conclude, the methodology presented here demonstrated the possibility of reconstructing the organization of myelinated fibers over large volumes in 3D at sub-micron resolution, enabling the study of the brain anatomy in both physiological and pathological conditions, thus offering a reliable method for integrated quantitative analyses. The versatility and simplicity of MAGIC will enable easy implementation of the technique in many laboratories, offering the possibility of using 3D investigation for routine analysis. We believe that MAGIC will help to have a more profound comprehension of the brain structure, bringing a significant impact on neuroscience.

## Methods

### Specimen collection

The investigated brain sections were obtained from postmortem brains from different species: mouse (control C57BL/6 and Reelin-deficient mouse model - RELN-/- Reeler, male, six months old), rat (Wistar, male, three months old), vervet monkey (African green monkey: Chlorocebus aethiops sabaeus, male, between one and two years old), and human (male, 87 years). The procedures for rodents were approved by the institutional animal welfare committee at Forschungszentrum Jülich GmbH, Germany, and were in accordance with the European Union guidelines for the use and care of laboratory animals. All methods were carried out in accordance with relevant guidelines and regulations. The vervet monkey tissue sample was acquired in the project “Postnatal development of cortical receptors and white matter tracts” (4R01MH092311-05) funded by the NIMH of the National Institutes of Health. The project was carried out in accordance with the UCLA Chancellor’s Animal Research Committee ARC #2011-135 and by the Wake Forest Institutional Animal Care and Use Committee IACUC #A11-219. The human brain was acquired in accordance with the ethics committee at the Medical Faculty of the University of Rostock, Germany #A2016-0083.

### MAGIC preparation protocol

Brains from different species (mouse, rat, vervet, and human) were fixed with 4% paraformaldehyde (PFA) solution at 4°C for several weeks (human brain: >3 months, vervet, rat, and mouse brains: 1–2 weeks). The brains were embedded first in a 10% glycerol, 2% DMSO, 4% formaldehyde solution at +4°C, then in a 20% glycerol, 2% DMSO, 4% formaldehyde solution at +4°C for mouse and rat brains 7 days in total, while for vervet and human brains ≥ 3 weeks. After treatment with 2% dimethyl sulfoxide for cryoprotection, brains were dipped in cooled isopentane (−50°C) for several minutes (mouse and rat brains: >5min, human and vervet brains: >30min). The frozen brains were cut with a cryostat microtome (Leica Microsystems, Germany) at a temperature of -30°C into sections of approximately 60 μm thickness. Brains were cut along one of three mutually orthogonal, anatomical planes: coronal, horizontal, or sagittal. Finally, the brain sections were incubated in a Phosphate Buffer Saline solution (PBS) 0.01M at room temperature (RT) for one month for mouse and rat brains and three months for vervet and human brains. Before imaging, the sections were mounted with PBS and coverslipped. For myelin characterization, the labeling was performed using the FluoroMyelin™ red dye (Thermo Fisher Scientific, cat. num. F34652): mouse brain sections were incubated in a solution of 1:300 FluoroMyelin™ in PBS for 20 minutes at room temperature. Then sections were rinsed 3 times for 10 minutes each with PBS.

### Fluorescence microscopy imaging

Fluorescence images were obtained using a commercial confocal microscope (Nikon Eclipse TE300 C2 LSCM, Nikon, Japan) equipped with a Nikon 60× or 100× immersion oil objective (Apo Plan, NA 1.4), and a custom-made two-photon fluorescence microscope (TPFM) at room temperature. Briefly, a mode-locked Ti: Sapphire laser (Chameleon, 120 fs pulse width, 90 MHz repetition rate, Coherent, CA) operating at 800 nm was coupled into a custom-made scanning system based on a pair of galvanometric mirrors (LSKGG4/M, Thorlabs, USA). The laser was focused onto the specimen by a refractive index tunable 25x objective lens (LD LCI Plan-Apochromat 25X/0.8 Imm Corr DIC M27, Zeiss, Germany) set either to glycerol or water. The imaged field of view was of 450 × 450 μm^2^, the resolution employed was 0.44 × 0.44 μm^2^ or 1.75 × 1.75 μm^2^. The system was equipped with a closed-loop XY stage (U-780 PILine® XY Stage System, Physik Instrumente, Germany) for the radial displacement of the sample and with a closed-loop piezoelectric stage (ND72Z2LAQ PIFOC objective scanning system, 2 mm travel range, Physik Instrumente, Germany) for the displacement of the objective along the z-axis. The fluorescence signal was collected by an independent GaAsP photomultiplier module (H7422, Hamamatsu Photonics, NJ). Emission filters of 482/35 nm and 618/50 nm were used for fibers and cell body detection, respectively.

### Mesoscale reconstruction

To perform mesoscale reconstruction of the samples, the volume of interest was acquired with TPFM performing z-stack imaging of adjacent regions using a custom LabView program (National Instruments). The 8-bit images (1024 × 1024 px or 256 × 256 px) produced were saved in .tiff format. Each stack was acquired with a depth equal to the thickness of the section (50 ± 10 μm) and with a z step of 1 or 2 μm between images. Each frame had a field of view of 450 × 450 μm^2^ and a pixel size of 0.44 × 0.44 μm^2^ or 1.75 × 1.75 μm^2^ for low-resolution reconstruction. The overlap of adjacent stacks was set as 40 μm. The stitching of all the acquired stacks was performed using ZetaStitcher (G. Mazzamuto, “ZetaStitcher: a software tool for high-resolution volumetric stitching” https://github.com/lens-biophotonics/ZetaStitcher). Low-resolution reconstructions were performed on the mouse coronal section, the rat sagittal section, and on the vervet section. High-resolution imaging was performed on the human hippocampus section and on the Reeler and control coronal section.

### Photon counting and Fluorescence Lifetime Imaging Microscopy (FLIM) measurements

FLIM measurements were performed on different mouse brain sections (N=15) during the three conditions of the protocol (samples: PFA N=4, Gly N=6, and MAGIC N=5) using a custom-made multimodal setup^16^. The collected fluorescence was sent to a high-speed PMT for photon counting PMH-100 (Becker-Hickl GmbH, Berlin, Germany) and then processed by a single-photon counting FLIM board SPC-730 (Becker-Hickl GmbH) for time-resolved analysis. Analysis of the obtained FLIM images was performed using the software SPC Image 4.9.7 (Becker-Hickl GmbH, Berlin, Germany), fitting the fluorescence decay data with a double-exponential decay function. From each image, we selected ROIs (10 for PFA, 12 for Gly, 8 for MAGIC) corresponding to the myelin sheath and the surrounding brain tissue in order to analyze the distribution of their lifetime values separately. The photon-counting obtained from the same samples were also used to evaluate the fluorescence efficiency of the MAGIC protocol. We measured the fluorescence intensity at different excitation wavelengths (from 740 nm to 880 nm), and we normalized it to the square of the respective laser power. For each condition, we selected the maximum amount of 240 μm^2^ ROIs detectable from the sample: 43 and 56 for PFA tissue and fibers respectively, 49 and 38 for Gly, 68 and 62 for MAGIC. Contrast evaluation was performed by dividing the fluorescence intensity detected from myelin fibers by that of the surrounding tissue.

### Fluorescence spectral measurements

The multimodal microscope, set at an excitation wavelength of 800 nm, was used to perform the spectral analysis measurements. Autofluorescence signals were collected during the three conditions of the protocol (N=1 sample for each step). The collected fluorescence signal was coupled to a multimode optical fiber by means of a 10× objective lens (Nikon, Tokyo, Japan) and detected in the 420-620 nm range using a multispectral detector PML-Spec (Becker-Hickl GmbH, Berlin, Germany) with 16 spectral channels. Each channel recorded a fluorescence intensity image in 90 s, from which we selected regions of interest (ROIs) corresponding to the myelin sheath (N=3) and the surrounding brain tissue (N=3) in order to analyze the intensity of their fluorescence emission separately.

### Raman measurements

We used a commercial Raman microscope (XploRA INV, Horiba, Kyoto, Japan) with λEX = 532 nm for collecting the Raman spectra of mouse brain tissues during the three conditions of the protocol on both the myelin sheath and the surrounding tissue (N=4 samples for each step). For each acquisition, we used a 60× objective (Nikon, Tokyo, Japan) for scanning a 10-µm-area while recording the Raman signal between 400 and 1750 cm^-1^ with 1800 lines/mm grating: each measurement lasted 30 s. The recorded spectra were processed to remove the fluorescence signal through an automated iterative routine (Vancouver Raman Algorithm). Each resulting Raman spectrum was normalized to its maximum intensity. Then, in order to evaluate the glycerol content within the examined tissue areas, we performed a spectral projection of all Raman spectra along with three major bands (550, 850 and 1465 cm^-1^) of glycerol; in particular, we calculated the scalar product between the spectra and the Raman peaks recorded from the glycerolized mounting medium solution. From that, for each spectrum, we obtained a score (“glycerol score”) equal to 1 for glycerolized tissues and <1 otherwise.

### Data analysis

Graphs and statistical analyses were done with OriginPro 9.0 (OriginLab Corporation) and www.socscistatistics.com. Mean and standard errors are displayed for each chart. Statistical analyses were performed using a one-tailed two-sample t-test. For the mean lifetime measurement the p-value (p), Cohen’s d (d), degrees of freedom (DF), and mean difference (m) ± confidence interval (CI) at 95% are: PFA fiber vs PFA tissue: p = 0.00028, d = 1.872, DF = 18, m+-95% CI = 98 ± 49; Gly fiber vs Gly tissue: p = 0.00041, d = 1.653, DF = 20, m+-95% CI = 109 ± 58; Magic fiber vs Magic tissue: p = 0.00009, d = 2.493, DF = 14, m+-95% CI = 150 ± 64. For the Magic fiber vs Magic tissue glycerol-score evaluation: p = 0.0173, d = 1.923, DF = 6, m+-95% CI = 0.06 ± 0.05. Stacks and 3D stitched volume renderings and videos were obtained using both Fiji (http://fiji.sc/Fiji) and Amira 5.3 (Visage Imaging).

### Structure tensor analysis (STA) evaluation

Following the preprocessing operations detailed in the Supplementary information, a structure tensor analysis (STA) was conducted at 5 µm spatial resolution on the whole mesoscale reconstruction of the hippocampus and separately on the selected ROIs (Figure 2 and Supplementary Fig.6) for estimating the local brain tissue orientation. To this end, we employed a custom STA tool developed by our laboratory in the framework of the European Human Brain Project. The source code of the present tool, written in Python3, can be accessed at: https://github.com/lens-biophotonics/st_fibre_analysis_hbp. In order to reject background and spurious dark regions, retaining only the contribution of brain structures, and improve the reliability of the obtained vector fields, a threshold of 85% non-zero voxels was imposed beforehand on each 5-μm macro-voxel to be characterized.

In detail, local gradient-square tensors were first computed as the outer product of the image gradient ∇*I*

with itself:

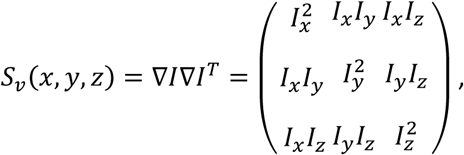

where *I*_*x*_, *I*_*y*_ and *I*_*z*_ respectively denote the local first-order spatial derivatives along the x, y, and z axes. Tensor elements estimated voxel-wise were then averaged over 5-µm local neighborhoods after isotropic smoothing by means of Gaussian kernels 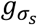 with standard deviation *σ*_*s*_ = 3 pixel.

3D tissue orientation maps were finally derived from the local directions of minimal gray level change, i.e. the eigenvector of the averaged structure tensor 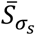 associated with the lowest eigenvalue.

### Orientation Distribution Functions (ODF) calculation

Fiber orientation distribution functions (ODFs) were used to characterize a given distribution of 3D orientations, i.e., nerve fibers. To calculate the ODF of a given number of K orientation vectors, the individual spherical harmonic (SH) coefficients c_lm_ of the ODF were estimated as 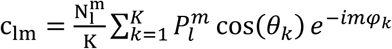 with 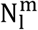 as the normalization coefficient, 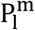 as Legendre polynomes, φ as azimuthal, and θ as the polar angle. This calculation was applied to the orientations obtained in a super-voxel consisting of n_x_ × n_y_ × n_z_voxels. In particular, to represent the ODFs of figure 2q and 2r a super voxel of 16 × 16 x 8 vectors, corresponding to a cube of 80 × 80 x 40 µm, was selected; while to obtain the total ODF of the full reconstruction and of the selected ROIs (represented in figure 2s), all the vectors present in the section were considered. The visualization of the ODFs was done with the open-source tool mrview from MRtrix3^17^. To avoid boundary artefacts, only super-voxels containing at least 1/3 of the total evaluated orientations were shown. The size of the ODFs was scaled by a factor of 2 to improve their visibility. Next, the MRtrix3 sh2peaks tool was applied to the obtained SH images in order to extract the Cartesian components of the three principal ODF lobes and, then, determine their amplitude (Euclidean norm). At this stage, amplitude values below 50% of the largest peak of the ODF were discarded, with the aim to exclude minor maxima related to noise from further consideration.

## Acknowledgments

We thank Markus Cremer, INM-1, Forschungszentrum Jülich, Germany for brain tissue sectioning and mounting. The research leading to these results has received funding from the European Union’s Horizon 2020 Framework Programme for Research and Innovation under the Specific Grant Agreement No. 785907 (Human Brain Project SGA2) and No. 945539 (Human Brain Project SGA3). This research has also been supported by the Massachusetts General Hospital (The General Hospital Corporation), Athinoula A. Martinos Center, The National Institute of Mental Health (NIMH) through the BRAIN Initiative Cell Census Network under award number 1U01MH117023-01, by the Italian Ministry for Education, University, and Research in the framework of the Eurobioimaging Italian Nodes (ESFRI research infrastructure) - Advanced Light Microscopy Italian Node, and by “Ente Cassa di Risparmio di Firenze” (private foundation) n.24135. The content is solely the responsibility of the authors and does not necessarily represent the official views of the National Institutes of Health.

## Author contributions

I.C. developed MAGIC, designed all the experiments, performed sample preparation, and conducted TPFM imaging. I.C and E.B performed the validation experiments. E.B and R.C analyzed the validation data. G.M developed the ZetaStitcher software and performed the mesoscopic reconstruction. F.G wrote the STA software. M.S evaluated the mesoscopic reconstruction with the STA software. F.M, M.M, and M.A effectuated the ODFs analysis. K.A and M.A provided the tissue specimens and contributed to the concept of the study. I.C, L.S, M.A, K.A, and F.S.P. supervised the project. I.C. made the figures and wrote the paper with inputs from all authors.

## Data availability

All data supporting the findings of this study are included in figures and videos as representative images or data points in the plots. Additional images other than the representative images are available from the corresponding author upon reasonable request.

## Code availability

MRtrix3 is an open source tool; ZetaStitcher and STA codes are open source and available on GitHub at URLs provided in the Methods. The other custom code used in this study are available from the corresponding author upon reasonable request.

