## Supplementary Information for "MAGIC: A label-free fluorescence method for 3D high-resolution reconstruction of myelinated fibers in large volumes"

### 1. Raman measurements and analysis

As reported in the main text, we used Raman spectroscopy for probing the glycerol content within myelinated fibers and the surrounding tissue before and after deglycerolization. In order to do so, we first recorded the Raman spectrum of the medium solution used for preparing and mounting glycerolized samples (20% glycerol, 2% DMSO, 4% formaldehyde). Supplementary figure 1a and 1b show the comparison between our measurement and the known spectra of glycerol and DMSO, respectively (spectra obtained from https://sdbs.db.aist.go.jp/sdbs/cgi-bin/cre_index.cgi). The main glycerol peaks can be easily identified in the recorded spectrum, while no major contribution can be attributed to the presence of DMSO. Then, we selected three major glycerol Raman bands (550, 850 and 1465 cm$^{-1}$) to be used for spectral projection: in fact, the "glycerol score" was calculated as the scalar product between each recorded spectrum and these three bands of the medium spectrum. Hence, a score = 1 means perfect match with the spectral signatures of glycerol, while a score < 1 implies lower abundance (or no presence at all) of that compound.

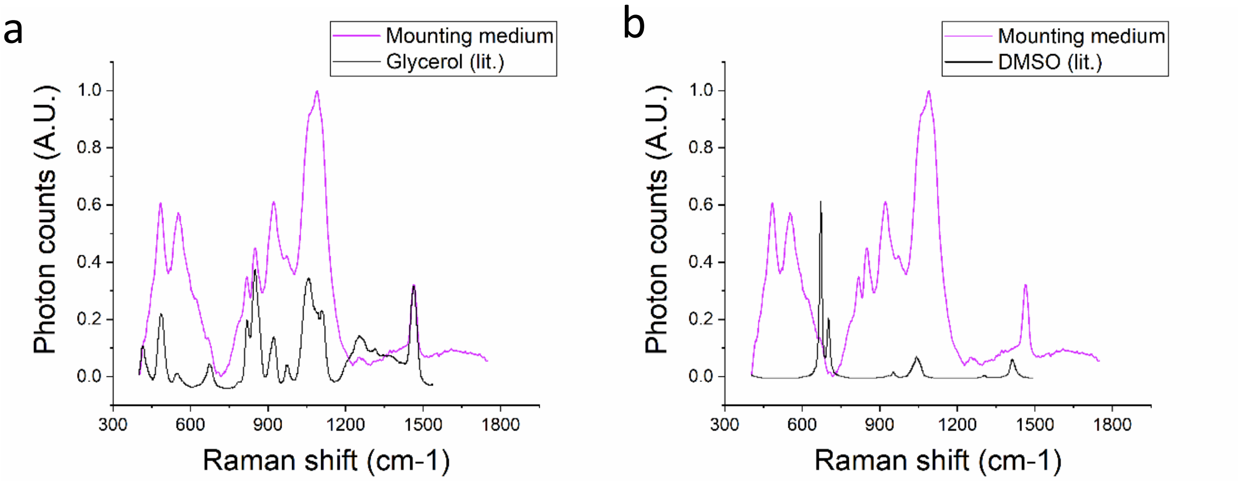


**Supplementary Figure 1:** (a) Overlaying Raman spectra of mounting medium (magenta) and literature glycerol spectra (black). (b) Overlaying Raman spectra of mounting medium (magenta) and literature DMSO spectra (black).

### 2. Compatibility with one photon imaging

The MAGIC protocol was developed to help the study of myelinated fibers with fluorescence microscopy techniques. In order to demonstrate the versatility of the protocol we performed acquisitions with a conventional confocal microscope at different wavelengths (Supplementary figure 2) and on different species (Supplementary figure 3).

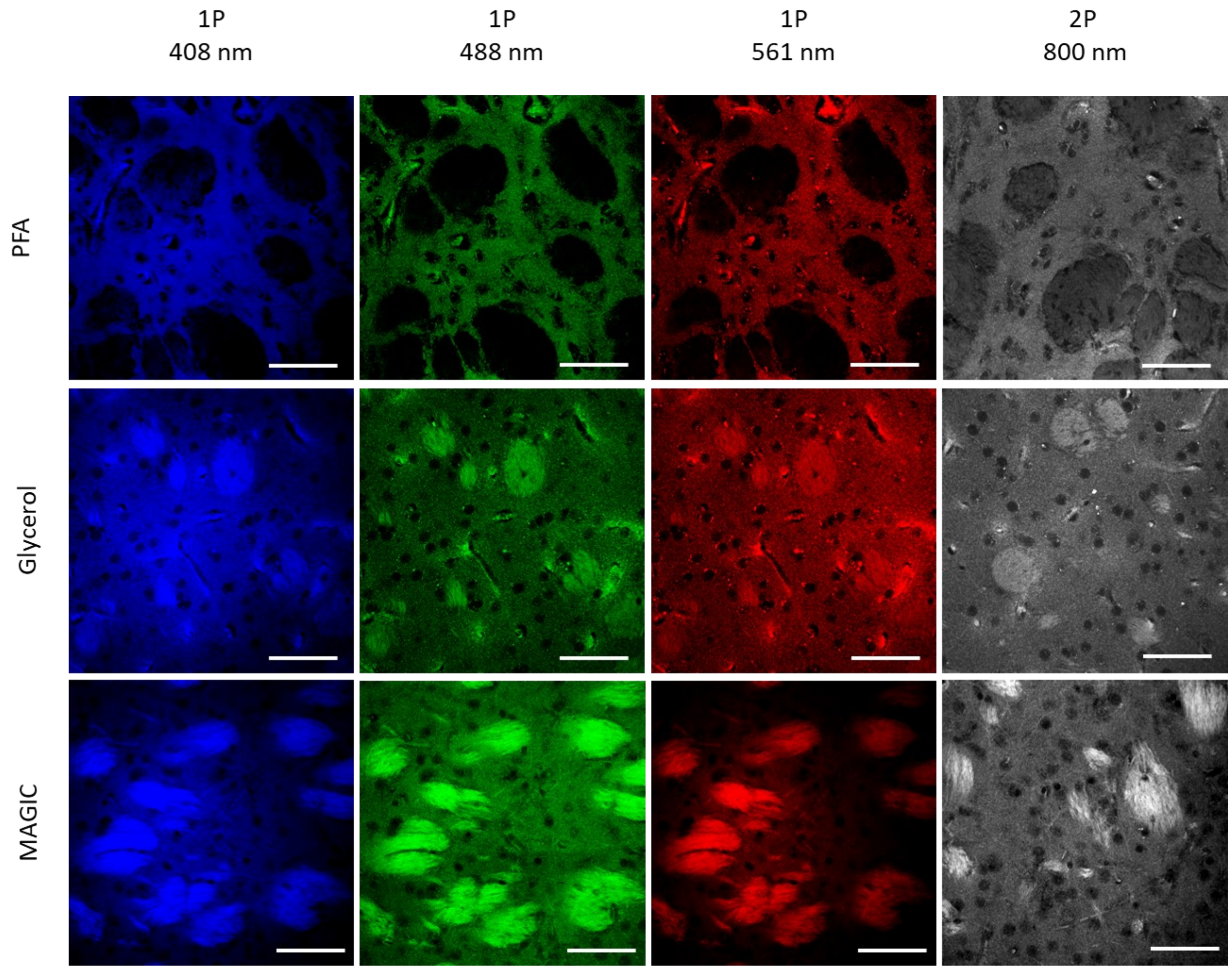


**Supplementary Figure 2:** Representative images of mouse brain sections during the three subsequent steps of the MAGIC protocol: fixation (PFA), glycerolization (Gly), and washing (MAGIC) acquired at different excitation wavelengths. One-photon (1P) excitation: 408, 488, 561nm; Two-photon (2P) excitation: 800nm. Scale bar = 50 µm.

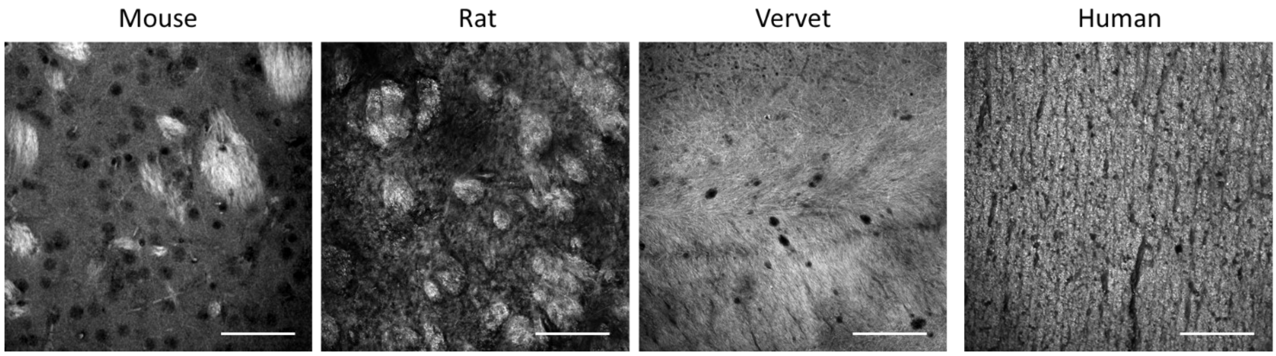


**Supplementary Figure 3:** Representative images of mouse, rat, vervet monkey, and human brain sections acquired with a confocal microscope using 488nm as excitation wavelength. Scale bar = 50 µm.

### 3. MAGIC protocol can provide details of myelin substructures

To establish the possibility of studying the different organization of the myelin sheaths surrounding axons, we used a high magnification objective (Nikon 100× immersion oil objective) to detect the autofluorescence signal coming from them. The high contrast obtained with the MAGIC methodology allows to measure both the inner and the outer diameters of the myelin sheaths surrounding axons as shown in supplementary figure 4.

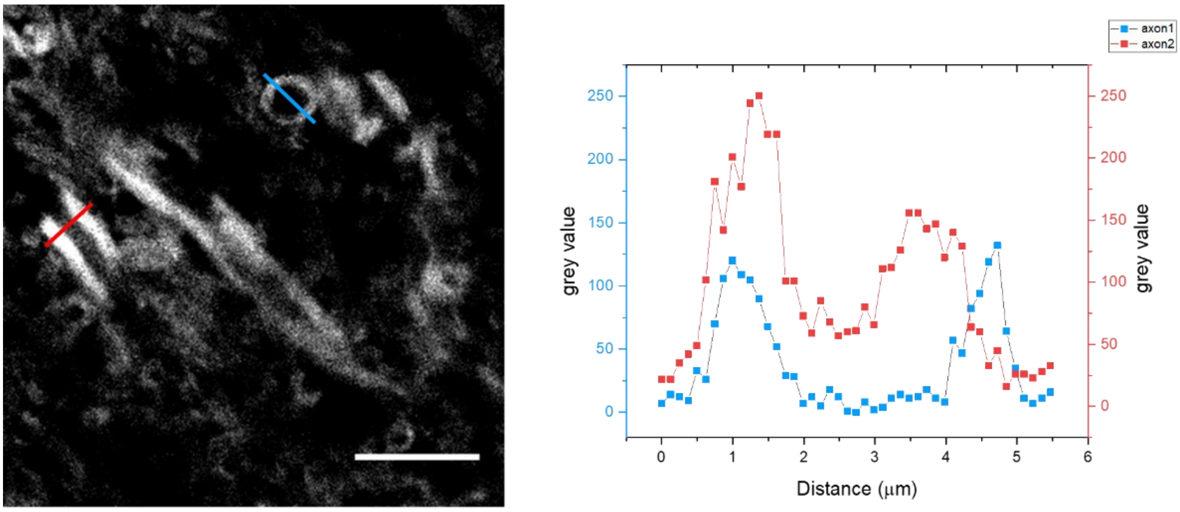


**Supplementary Figure 4:** Image obtained with a 100x objective ($\lambda_{exc}$=488nm), Scale bar = 10 μm. The graph shows the intensity profile of the lines corresponding to the two axons (blue and red) of the image.

### 4. MAGIC and Immunostaining

To test the compatibility of the MAGIC protocol with conventional immunofluorescence, human brain sections treated with MAGIC were stained with an anti-GFAP antibody. Astrocytes were successfully stained, representative images are shown in supplementary figure 5, and in supplementary videos 2 and 3. To perform the staining, the following protocol was applied. Brain sections previously treated with MAGIC protocol were permeabilized, incubating the sample for 45 min with PBS with 0.5% Triton X-100. After, samples were incubated with the primary antibody, an anti-GFAP antibody (abcam; ab7260) with a dilution of 1:200 in PBST 0.1% overnight at 4°C in the dark. The solution containing the primary antibody was removed and the sample was washed three times for 8 hours in PBST 0.1% solution. The day after, the sample was incubated with the secondary antibody, a donkey anti-rabbit IgG antibody conjugated with an alexa fluor 568 with a dilution of 1:200 (abcam, ab175470) in PBST 0.1% for 2 h at room temperature in the dark. Secondary antibody was removed, and samples were washed three times with PBST 0.1% for 2h each in the dark at @RT. Finally, sample were washed two times with PBS for 10 min each at @RT, coverglass was mounted on the section with transparent nailpolish, and TPFM imaging was performed.

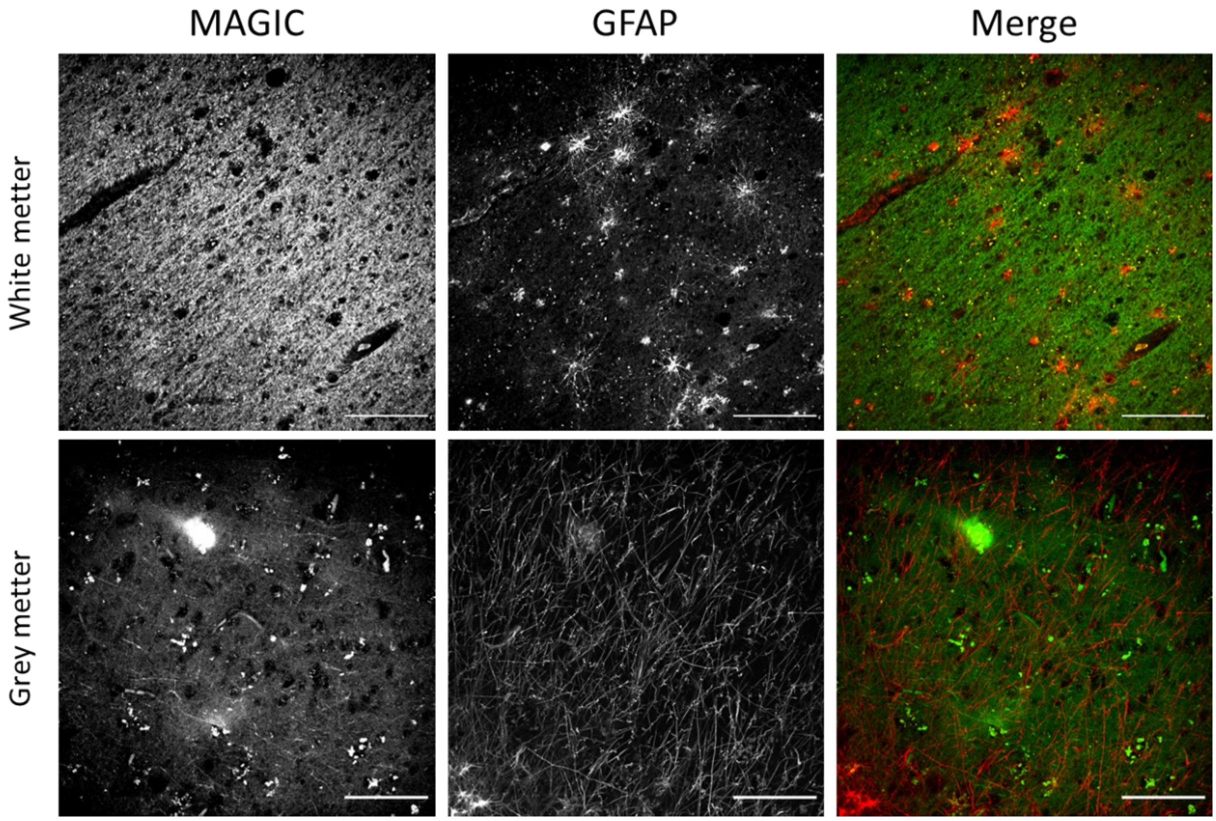


**Supplementary Figure 5:** Images showing fibers (in green; MAGIC) and astrocytes (in red; anti-GFAP antibody) in both white and grey matter of a human brain section. Scale bar = 100 μm.

### 5.Fiber analysis pre-processing

Reeler mouse (Reelin deficient - RELN-/- Reeler) is a well-known animal model for several neurological and neurodegenerative disorders. Here we applied our method to a brain section (stereotactic coordinate: -2.18 mm Bregma; 1.62 mm Interaural) from one control mouse (Control) and a reeler mouse (Reeler) to observe the different structural organization in various areas of the right hippocampal region. In order to perform the analysis, a manual segmentation of the areas of interest was performed using Fiji (http://fiji.sc/Fiji). The supplementary figure 6 shows the different masks superimposed on the maximum intensity projection (MIP) of the mosaic acquired with the TPFM.

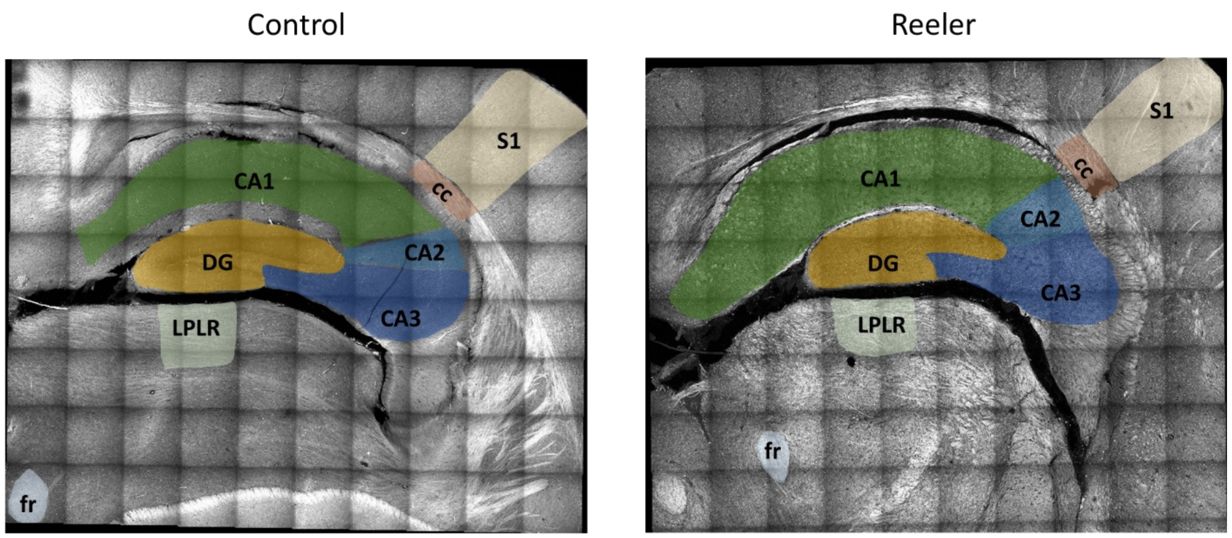


**Supplementary Figure 6:** Masks of different areas superimposed to the MIP of the mosaic of the right hippocampal region of the Control and the Reeler mouse sections. Acronym list = S1: Primary Somatosensory Cortex; cc: Corpus Callosum; CA1, CA2, CA3: field CA1, CA2, CA3 of hippocampus; DG: Dentate Gyrus ; LPLR: Lateral Posterior Thalamic Nucleus; fr: Fasciculus Retroflexus

### 6. Structure tensor analysis and orientation distribution functions evaluation

In order to compensate for the relative inclination of the brain sections with respect to the coronal image plane, the whole mesoscale stack mosaic of the hippocampus and its segmented ROIs were initially rotated along the frontal and longitudinal axes using Fiji. Next, image blurring was applied in the coronal plane so as to uniform the in-plane $FWHM_{XY}$ of the optical system PSF ($FWHM_{XY}$ = 0.692 µm) to the sagittal $FWHM_{Z}$ (2.612 µm).

After these preprocessing operations, stack volumes were virtually decomposed into macro-voxels of (12, 12, 5) pixel size for obtaining a 5-µm tissue analysis resolution, and then downscaled within the image plane in order to set an isotropic spatial sampling period of 1 µm. As mentioned in the main text, a threshold of 85% non-zero voxels was applied to the separate 5-µm image sub-blocks with the aim to exclude the stack background and local regions associated with a scarce proportion of brain tissue data. In practice, this macro-voxel selection step was instrumental in improving the accuracy of the resulting orientation estimates in boundary macro-voxels and in rejecting the contribution of spurious dark regions.

Afterwards, for each accepted macro-voxel, local (voxel-size) image gradient-structure tensors $S(x,y,z)$ were first computed as:

$$S(x,y,z) = \nabla I \nabla I^T * g_{\sigma_s} = \begin{pmatrix} I_x^2 * g_x & I_x I_y * g_y & I_x I_z * g_z \\ I_y I_x * g_x & I_y^2 * g_y & I_y I_z * g_z \\ I_z I_x * g_x & I_z I_y * g_y & I_z^2 * g_z \end{pmatrix},$$

where the first-order spatial derivates $I_x$ , $I_y$ and $I_z$ were estimated using second-order accurate central differences, whereas $g_x$, $g_y$ and $g_z$ represent Gaussian smoothing filters with standard deviation $\sigma_s = 3$ pixel.

Local $S(x,y,z)$ elements were then averaged within each of the distinct 5-µm macro-voxels and the spectral decomposition reported below was performed:

$$\bar{S} = \begin{pmatrix} \bar{S}_{xx} & \bar{S}_{yx} & \bar{S}_{zx} \\ \bar{S}_{yx} & \bar{S}_{yy} & \bar{S}_{zy} \\ \bar{S}_{zx} & \bar{S}_{yz} & \bar{S}_{zz} \end{pmatrix} = \begin{pmatrix} | & | & | \\ v_1 & v_2 & v_3 \\ | & | & | \end{pmatrix} \begin{pmatrix} \lambda_1 & 0 & 0 \\ 0 & \lambda_2 & 0 \\ 0 & 0 & \lambda_3 \end{pmatrix} \begin{pmatrix} | & | & | \\ v_1 & v_2 & v_3 \\ | & | & | \end{pmatrix}^{-1}$$

e-vectors e-values

Finally, an eigenvalue analysis was conducted on the resulting decomposition for identifying the eigenvectors $v_i$ associated with the lowest eigenvalue $\lambda_i < \lambda_j < \lambda_k$ $(i, j, k \in [1,2,3]$ , axes) thus generating 3D tissue orientation maps corresponding to the directions of minimal gray level change at 5-um resolution. Orientation Distribution Functions ODFs evaluation was performed on the resulted vectors as described in the main text.

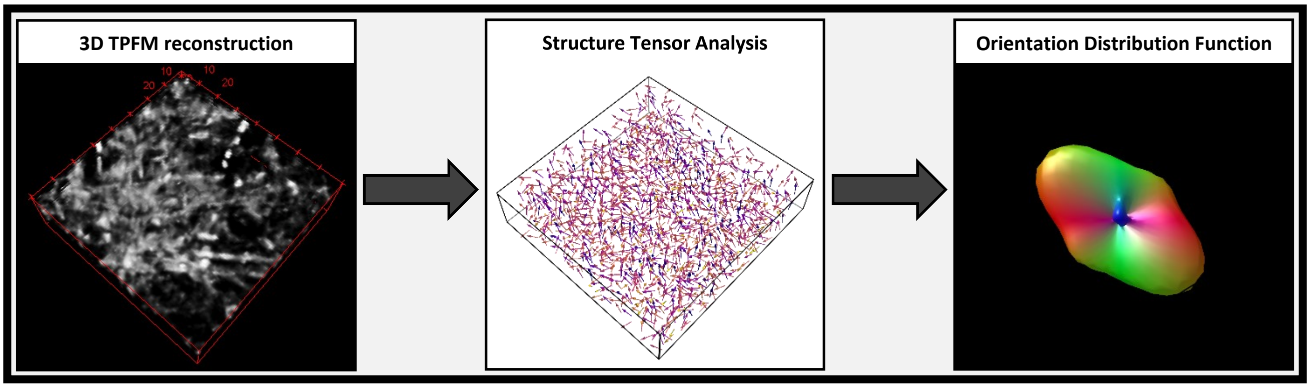


**Supplementary Figure 7:** block scheme representing the evaluation conducted on the TPFM images.

| ROI | CONTROL MOUSE | | | | | | | REELER MOUSE | | | | | | |
|---|---|---|---|---|---|---|---|---|---|---|---|---|---|---|
| | ODF Lobes | $a_1$ | Direction | $a_2$ | Direction | $a_3$ | Direction | ODF Lobes | $a_1$ | Direction | $a_2$ | Direction | $a_3$ | Direction |
| Full | 3 | 0.14 | In-plane | 0.14 | In-plane | 0.09 | Out-of-plane | 3 | 0.16 | Out-of-plane | 0.11 | In-plane | 0.11 | In-plane |
| cc | 1 | 0.35 | In-plane | | | | | 1 | 0.26 | In-plane | | | | |
| S1 | 2 | 0.20 | In-plane | 0.12 | In-plane | | | 3 | 0.17 | In-plane | 0.11 | In-plane | 0.11 | Out-of-plane |
| DG | 1 | 0.16 | In-plane | | | | | 1 | 0.19 | Out-of-plane | | | | |
| CA1 | 1 | 0.23 | In-plane | | | | | 2 | 0.18 | Out-of-plane | 0.09 | In-plane | | |
| CA2 | 1 | 0.18 | In-plane | | | | | 1 | 0.24 | Out-of-plane | | | | |
| CA3 | 2 | 0.15 | In-plane | 0.08 | Out-of-plane | | | 1 | 0.20 | Out-of-plane | | | | |
| fr | 1 | 0.23 | In-plane | | | | | 2 | 0.21 | Out-of-plane | 0.18 | In-plane | | |
| LPLR | 1 | 0.30 | In-plane | | | | | 2 | 0.18 | Out-of-plane | 0.16 | In-plane | | |

**Supplementary table 1**: Amplitude (a1, a2, a3) and main orientation (direction) of the significant ODF components are shown. Table cell is gray if the peak amplitude is <50% of the primary peak. Acronyms list = Full: full field of view; S1: Primary Somatosensory Cortex; cc: Corpus Callosum; CA1, CA2, CA3: field CA1, CA2, CA3 of hippocampus; DG: Dentate Gyrus; fr: Fasciculus Retroflexus; LPLR: Lateral Posterior Thalamic Nucleus.

### Supplementary videos legend:

**Video1:** Navigation in the 3D rendering of myelinated fibers after MAGIC. Imaging performed with TPFM, volume of 200 x 200 x 40 $\mu m^3$.

**Video2:** 3D rendering navigation of a human brain white matter stack labeled with anti-GFAP after MAGIC. Imaging performed with TPFM: myelinated fibers in green and astocytes in red. Volume of 450 x 450 x 60 $\mu m^3$.

**Video3:** Rotation of a 150 x 150 x 60 $\mu m^3$ ROI of the video2 showing a single astrocyte. Imaging performed with TPFM: myelinated fibers in green and astocytes in red.

**Video4:** 3D rendering navigation of a representative stack of the human hippocampus after MAGIC. Imaging performed with TPFM: myelinated fibers in green and cell bodies in red due to lipofuscin autofluorescence. Volume of 450 x 450 x 60 $\mu m^3$.
